# Magnetoencephalography can reveal deep brain network activities linked to memory processes

**DOI:** 10.1101/2022.02.28.482228

**Authors:** Víctor J. López-Madrona, Samuel Medina Villalon, Jean-Michel Badier, Agnès Trébuchon, Velmurugan Jayabal, Fabrice Bartolomei, Romain Carron, Andrei Barborica, Serge Vulliémoz, F. Xavier Alario, Christian G. Bénar

**Affiliations:** Aix Marseille Univ, INSERM, INS, Inst Neurosci Syst, Marseille, 13005, France; APHM, Timone Hospital, Epileptology and cerebral rhythmology, Marseille, 13005, France; APHM, Timone Hospital, Functional and stereotactic neurosurgery, Marseille 13005, France; Physics Department, University of Bucharest, 405 Atomistilor Street, Bucharest, Romania; EEG and Epilepsy Unit, University Hospitals and Faculty of Medicine Geneva, Rue Gabrielle-Perret-Gentil 4, 1205 Geneva, Switzerland; Aix-Marseille Université, CNRS, LPC, Marseille, France

## Abstract

Recording from deep neural structures such as hippocampus non-invasively and yet with high temporal resolution remains a major challenge for human neuroscience. Although it has been proposed that deep neuronal activity might be recordable during cognitive tasks using magnetoencephalography (MEG), this remains to be demonstrated as the contribution of deep structures to MEG recordings may be too small to be detected or might be eclipsed by the activity of large-scale neocortical networks. In the present study, we disentangled mesial activity and large-scale networks from the MEG signals thanks to blind source separation (BSS). We then validated the MEG BSS components using intracerebral EEG signals recorded simultaneously in patients during their presurgical evaluation of epilepsy. In the MEG signals obtained during a memory task involving the recognition of old and new images, we identified with BSS a putative mesial component, which was present in all patients and all control subjects. The time course of the component selectively correlated with SEEG signals recorded from hippocampus and rhinal cortex, thus confirming its mesial origin. This finding complements previous studies with epileptic activity and opens new possibilities for using MEG to study deep brain structures in cognition and in brain disorders.

## INTRODUCTION

Magnetoencephalography (MEG) is a non-invasive technique that measures the electromagnetic fields generated by the brain with a millisecond time scale. MEG is widely used to analyze cortical activity in a variety of scientific and clinical settings (Baillet, 2017; Bartolomei et al., 2006; Hämäläinen et al., 1993; Wacongne et al., 2011). In most of these applications, a challenging inverse problem needs to be solved in order to determine the neuronal sources generating the recorded MEG signal (Fokas et al., 2004; Friederici et al., 2000; Hillebrand et al., 2005). Neuronal sources located relatively near the sensors (i.e. at the cortical surface) are notoriously easier to identify than sources located more distally (Huotilainen et al., 1998; Mosher et al., 1993). Sources located in deep neuronal structures such as mesial structures provide the more challenging case, and the reliability of their identification is a matter of ongoing debates (Bénar et al., 2021; Pu et al., 2018; Ruzich et al., 2019). Biophysical models have suggested how, under certain conditions, it might be possible to identify the magnetic activity originating from deep structures and to separate the activity of various closed-field sources (Attal et al., 2012; Meyer et al., 2017; Stephen et al., 2005). Some empirical studies have reported deep MEG sources taken to reflect hippocampal activity (Barry et al., 2019; Costers et al., 2020; Fellner et al., 2019; Guderian et al., 2009; Taylor et al., 2011; Xu et al., 2020). This interpretation hinged primarily on the link between the activity of interest and the behavioral performance in a cognitive task, but the studies lacked a neuronal ground truth that could confirm the origin of the signals.

Intracerebral recordings, in the form of Stereo-ElectroEncephaloGraphy (SEEG, Bancaud et al., 1970), give an unrivalled opportunity for measuring directly time-resolved neuronal activities in human deep brain structures. This invasive technique is commonly used as part of the pre-surgical diagnosis procedure in drug-resistant epilepsy to delineate the epileptogenic network (Bartolomei et al., 2017). It has been successfully employed in cognitive studies, allowing to track both evoked and oscillatory activities (Barbeau et al., 2017, 2008; Despouy et al., 2020; Hagen et al., 2020; Jonas et al., 2016; Lachaux et al., 2005; Nelson et al., 2017; Tallon-Baudry et al., 2005). Because the SEEG signal is recorded locally within the implanted structures, it can be taken as a unique ground truth to which non-invasive measures can be confronted to, although recent studies have suggested that recording EEG or MEG responses to intracranial stimulation may represent a ground truth for source localization algorithms (Mikulan et al., 2020; Parmigiani et al., 2021). Still, in order to optimize the measure of correspondence between MEG and intracranial data, their recordings should be simultaneous (Rampp et al., 2010; Shigeto et al., 2002; Sutherling et al., 2001). Simultaneous recordings are the only ones that allow computing time-resolved correlations between depth and surface signals (Boran et al., 2020; Crespo-García et al., 2016; Dalal et al., 2009; Dubarry et al., 2014; Korczyn et al., 2013; Pizzo et al., 2019), thus ensuring that the very same activity is measured on both recordings. Studies based on simultaneous recordings present the strongest evidence supporting the detectability of deep activity (Crespo-García et al., 2016; Dalal et al., 2009; Korczyn et al., 2013), with a strong correlation between contacts placed in the hippocampus and the MEG signals (Dalal et al., 2009).

Surface signals are non-invasive but capture a complex mixture of signals, where hippocampal activity proper might be present but hidden. Spatial filtering, either within a source localization framework or with blind source separation (BSS), can help retrieving activity from deep structures, disentangling it from more superficial signals that present higher amplitudes on MEG (Attal et al., 2007; Attal and Schwartz, 2013; Dubarry et al., 2014; Oswal et al., 2016; Pizzo et al., 2019). Importantly, BSS exploits the sparsity of the brain sources (Daubechies et al., 2009), which has been pointed out as a key element for the identification of deep activity (Krishnaswamy et al., 2017). In prior work from our laboratory involving patients with temporal lobe epilepsy, we used independent component analysis, a kind of BSS, to show that interictal spikes generated in the hippocampus and amygdala were indeed detectable with MEG (Pizzo et al., 2019). However, the amplitude of the epileptic spikes is much higher than the spontaneous non-pathological activity, and therefore easier to detect from the surface. Using a cognitive paradigm, Dalal and collaborators have shown zero-lag correlation for theta activity in the hippocampus (Korczyn et al., 2013), although the possibility of zero-lag phase synchrony with a third structure could not be ruled out in that dataset.

Here we sought to firmly establish the link between MEG and SEEG signals evoked by a cognitive task. We used a well-tested experimental paradigm that activates mesial structures during short-term picture memorization and recognition. We acquired simultaneously MEG and SEEG and used BSS to quantify the single-trial correlation between BSS-MEG components and intracerebral data across a total of six patients. Additionally, we recorded the MEG signals from six healthy volunteers as a control. Based on the available biophysical models, we hypothesized that the activation of mesial structures during memory processes would be detected with MEG.

## METHODS

### Patients and records selection

We studied six patients (two females) by means of simultaneous MEG-SEEG recordings. Table 1 shows the clinical information for each patient. We also recorded and analyzed MEG data from six healthy volunteers (three females, mean age 31.7 years SD ± 5.5) performing the same protocol. This research has been approved by the relevant Institutional Review Board (Comité de Protection des Personnes, Sud-Méditerranée I, ID-RCB 2012-A00644–39). All participants signed a written informed consent form regarding this research.

**Table 1:**
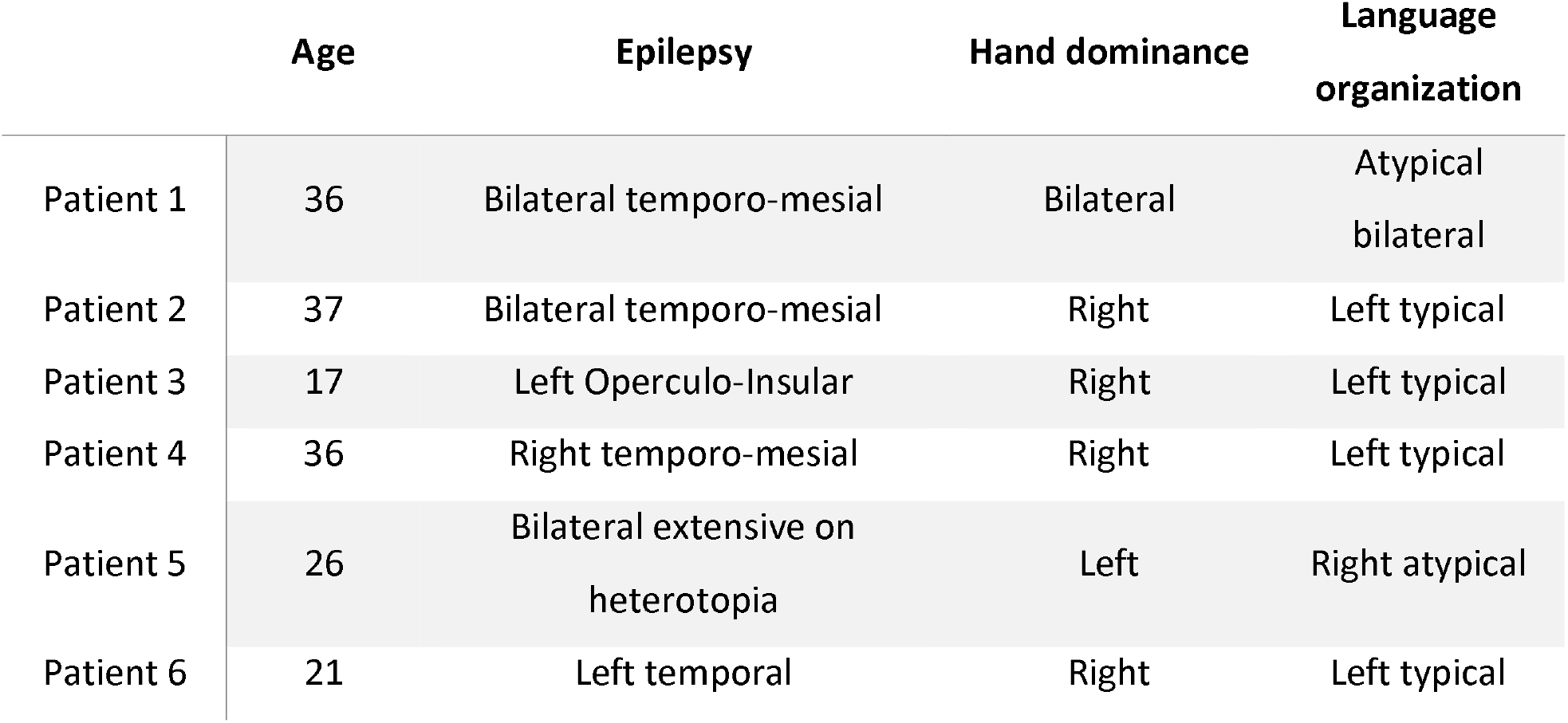
Clinical information of each patient.

|  | Age | Epilepsy | Hand dominance | Language organization |
| --- | --- | --- | --- | --- |
| Patient 1 | 36 | Bilateral temporo-mesial | Bilateral | Atypical bilateral |
| Patient 2 | 37 | Bilateral temporo-mesial | Right | Left typical |
| Patient 3 | 17 | Left Operculo-Insular | Right | Left typical |
| Patient 4 | 36 | Right temporo-mesial | Right | Left typical |
| Patient 5 | 26 | Bilateral extensive on heterotopia | Left | Right atypical |
| Patient 6 | 21 | Left temporal | Right | Left typical |

### Simultaneous SEEG-MEG recordings

The six patients were undergoing intracerebral stereotaxic EEG (SEEG) investigation for presurgical evaluation of focal drug-resistant epilepsies at the Epileptology and Cerebral Rhythmology Unit, APHM, Marseille, France. We acquired SEEG and MEG recordings simultaneously during 15 minutes of resting state (patient relaxed with eyes closed) and subsequently during a memory task (see below). The methodology for the simultaneous recordings is detailed in previous studies (Badier et al., 2017; Dubarry et al., 2014). MEG signals were acquired on a 4D Neuroimaging™ 3600 whole head system with 248 magnetometers at a sampling rate of 2034.51 Hz. We acquired between 161 and 223 SEEG contacts per patient (total contacts recorded: 1206; mean of 201 contacts, SD ± 23) at 2048 Hz of sampling rate, as well as EOG and ECG channels. The electrodes had a diameter of 0.8 mm, and contained 10 to 15 contacts, each being 2 mm long and separated from each other by 1.5 mm (Alcis, Besançon, France). The implantation of the intracranial SEEG was decided based on clinical hypotheses regarding the location of the epileptogenic zone. A description of the SEEG setup is detailed in Pizzo et al. (Pizzo et al., 2019).

### Memory protocol

Each block of the recognition memory task started with an encoding phase, during which 12 pictures were presented, one after the other, and the participant was asked to memorize them. Each picture was a simple colored drawing of a familiar item (e.g. a dog or a car) on a grey background; pictures were obtained from the standardized database Multipic (Duñabeitia et al., 2018). After a distracting video of one minute (silent excerpts from a documentary showing birds and landscapes), the recognition phase involved a larger set of pictures from the same source. Half of these pictures had been presented during the encoding phase, while the other 12 were new never-presented pictures. Participants were asked to press a button with the right hand if they recognized the image as having been presented earlier (“old”) during encoding, or a button with the left hand if the image was “new” to them. Stimuli presentation and response logging were controlled by the software E-prime 3.0 (Psychology Software Tools, Pittsburgh, PA).

Each trial started with a fixation cross presented in the center of the screen for 1000 ms, followed by the experimental picture, presented for 1000 ms in the encoding sub-block and for 1500 ms in the recognition sub-block. The subsequent inter-trial interval was fixed to 1000 ms. For each participant, a total of 7 blocks were programmed to be displayed consecutively, using different images.

We selected 24 × 7 = 168 images to be used as experimental materials from the database of Duñabeitia et al., 2018. They were selected as having high name agreement (above 90%), and a relatively short name (1, 2, or 3 syllables in French). To ensure that the observations were not driven by item-specific properties, different experimental lists were created for each participant. The items were separated into two matched groups of 84 items to serve as old and new, alternatively across patients. Across the “old” and “new” groups of items there were roughly equal numbers of natural and artefact stimuli, with matched visual complexities; their names in French were matched for name agreement, length in syllables, and (log) lexical frequency of use (normative data from Duñabeitia et al., 2018 or New et al., 2004). The 84-item groups were further broken down into 7 groups to be used in the different blocks, with items matched for visual complexity and (log) lexical frequency across the 7 groups. All matching across picture groups was performed with the MATCH utility (van Casteren and Davis, 2006). In the encoding phases, the 12 items were presented in a random order; in the recognition phases, the items were presented in a pseudo-random order, with the constraint that there were never more than 3 “old” or “new” items in a row.

### MEG and SEEG pre-processing

All data analysis was done using a combination of the in-house AnyWave software (Colombet et al., 2015; available at http://meg.univ-amu.fr/wiki/AnyWave), the Fieldtrip toolbox (Oostenveld et al., 2011), and custom-made Matlab scripts (The Mathworks Inc., Naticks, MA, USA). Each trial was epoched from 500 ms pre-stimulus to 1500 ms post-stimulus. After visual inspection, MEG and SEEG channels with noise or flat signal were removed, as well as all trials with artifacts in either the MEG or SEEG signal. Continuous data was bandpass filtered between 0.5 and 120 Hz (FIR filter) and two notch filters at 50 and 100 Hz were applied to remove line noise and its first harmonic. To remove eye blinks, movements and cardiac components on MEG data, independent component analysis (ICA) was computed on the cleaned data using the Infomax algorithm (Bell and Sejnowski, 1995) implemented in AnyWave. Before ICA, we performed a principal component analysis (PCA) to reduce the number of dimensions to 100. Based on the time course and the scalp topography, components related to cardiac activity or eye blinking were rejected (Jung et al., 2000).

### Separation of neuronal sources

To identify and separate the different brain sources that contribute to the MEG recordings, we used second-order blind identification (SOBI; Belouchrani et al., 1997, 1993; Tang et al., 2005a). SOBI takes advantage of the temporal correlation within sources; it finds an unmixing matrix by minimizing the sum-squared cross-correlation between one component at time t and another component at time t + s, across a set of time delays. This way, SOBI allows the identification of highly temporally correlated neuronal sources (Belouchrani et al., 1993).

For each patient, we concatenated all trials across all conditions, seeking one unmixing matrix per participant. We applied PCA on the preprocessed MEG data to reduce its dimensionality to 100. Then, we computed SOBI using the Fieldtrip algorithm with 100 time delays (~50 ms, Tang et al., 2005a). For comparison purposes, we repeated the analysis with 50 and 150 dimensions, without notable differences in the results. Although only some of the components were putative neuronal sources, we did not discard any of them at this point.

### Identification of components responding to the stimulus

For each SOBI and SEEG channel we checked if they had a significant event-related potentials (ERP) based on all triggers (old and new). To do so, for each channel we tested that each time point across trials was significantly different from zero with a t-test, obtaining a p-value and a t-value. Then, we used the local false discovery rate (LFDR; Benjamini and Heller, 2007) with a LFDR alpha of 0.2 (Pizzo et al., 2019). This resulted in a threshold on the t-values that takes into account multiple comparisons between samples and components. Briefly, LFDR identifies which values stand out from the noise, whose distribution is assumed to be Gaussian. In our case, given a distribution with all t-values, LFDR determines the threshold (one positive and another negative) from which the values are considered statistically significant. For example, we computed one LFDR for all SOBI components during recognition (“old” trials), obtaining a single threshold for all the datasets of a participant in that condition. To remove artifactual single points, we selected only those points during the first second after the stimulus and we imposed a minimum number of consecutive significant time samples (10 ms in this work). We repeated this process in each subject and condition, for SEEG and for SOBI data separately.

### Selection of deep SOBI-MEG components

First, we rejected all the components with a noisy topography. To determine putative deep SOBI components, we visually reviewed all the components with a significant response and compared their topographies across patients. We identified only one component with a robust topography that was similar across patients and that reflected a putative deep origin. The source and sink of the dipolar topography were far from each other, indicating an origin remote from the recording sensors. We selected this SOBI component as a putative deep SOBI-MEG.

### Differences between “old” and “new” conditions

We evaluated whether each SEEG and SOBI component with a significant ERP was also modulated as a function of the experimental conditions. For each patient and time point, we compared with a t-test defined across trials the amplitude of the ERPs between old and new conditions. Then, to correct for multiple comparisons, we computed LFDR on the t-values for each dataset (i.e., each patient), leading in a test for all SEEG channels and SOBI components for significant ERP differences.

### Depth-surface temporal correlation

We computed the zero-lag correlation (Matlab function *corrcoeff*) between the SEEG and SOBI components on their continuous time-series during stimuli presentation (around 8 minutes of recordings). The use of many time points implies that even very low values of correlation can be identified as significant. Here too, we used LFDR on the correlation values to determine which pairs of SEEG-SOBI correlations were statistically significant. To improve the distribution estimation in the LFDR analysis, we included all the SEEG and SOBI-MEG signals, not only those with a significant ERP. in. We applied a Fischer transformation on the correlation coefficients to approximate a Gaussian distribution (Pizzo et al., 2019). Then, we applied LFDR on all the correlation values for each patient, considering as significant those pairs with a correlation, in absolute value, higher that the LFDR threshold.

### Partial correlation between SOBI-MEG and SEEG across mesial structures

We determined the location of each SEEG contact using the software GARDEL, (available at https://meg.univ-amu.fr, Medina Villalon et al., 2018). This tool allows automatic electrode localization and contact labelling according to a given atlas. Here, we used the VEP atlas (Wang et al., 2020), based on Destrieux parcellation (Destrieux et al., 2010) and subdivided to correspond more to anatomical and functional clinical areas. We selected three regions located in the temporal lobe: anterior hippocampus, rhinal cortex, and middle temporal gyrus. We chose these structures because they were recorded in most of the patients (four out of five) and because they presented differences between the old and new conditions in all patients. For each region, we selected the contact with the highest correlation with the SOBI-MEG. To discriminate the contribution of each region to the SOBI-MEG, we applied a partial correlation analysis between the continuous data of the three SEEG contacts and the SOBI-MEG (Marrelec et al., 2006). For each pair of signals, the partial correlation aims to disentangle the information present only in both signals (direct correlation) from the activity that originates in other regions (indirect correlation).

To determine whether the values of partial correlation were significantly different from zero we followed a surrogate approach (Cohen, 2014). We used the continuous time-series (i.e. trials were concatenated) to generate the surrogates. For each patient we created a subset of surrogate data (N=1000) by randomly displacing in time the SEEG traces. Each SEEG signal was divided into two segments of random length and their order was switched. This way, the temporal relationship between signals is broken, but their autocorrelation is preserved (Pereda et al., 2005). We then computed the partial correlation between the surrogate SEEG and the SOBI-MEG. The surrogate results of each region approximate a normal distribution. Thus, the p value of significance can be computed as the distance (in units of standard deviation) from the original value to the mean of the surrogate distribution. The threshold of significance was set to p=0.025 for each patient and region.

### Source localization of SOBI-MEG

If SOBI correctly separates brain sources into different components, the obtained component topographies are expected to be dipolar (Delorme et al., 2012). Their localization can be estimated with an equivalent current dipole. In some cases, the SOBI components may correspond to a bilateral activation of homologous brain regions that can be modeled by symmetric dipoles (Bénar et al., 2021; Piazza et al., 2020). We assumed the latter for the SOBI-MEG, representing a bilateral activation of the mesial network. Thus, to localize the source of each SOBI-MEG topography, we applied a two-dipole fitting procedure that was symmetric with respect to the longitudinal plane. For the forward model, we used the shaped single shell approximation implemented in FieldTrip (Oostenveld et al., 2011), which is based in Nolte’s solution (Nolte, 2003). Each point of a grid within the brain volume was associated to a triplet of orthogonal dipoles. We compared the projection of the model composed by those triplets with the SOBI topography, computing the goodness of fit (GOF) at each location as one minus the ratio of the sum of squared difference between the SOBI map and the model divided by the sum of square of the SOBI map. A confidence interval was estimated by including all points between the maximum GOF minus the distance from the maximum GOF to 1 (Pizzo et al., 2019):

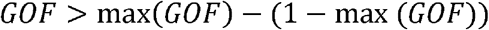

## RESULTS

### Identification of putative deep sources on MEG with SOBI

We performed simultaneous recordings of MEG and SEEG in six patients with focal drug-resistant epilepsy. To identify the different sources mixed on the MEG signals, we used SOBI on the MEG signals recorded during the task. A visual review revealed one SOBI-MEG component that was present in all patients, with similar topographies (Figure 1) and ERPs (Figure 2). We selected this component as a putative deep source. Its topography resembles a single dipole located far from the surface, with the positive and negative poles of the radial component of the magnetic field topography over the temporal lobes (Figure 1a). Its ERP presents a complex pattern (Figure 2a). It has the earliest response at 110 ms, which matches in time with the N110 visual component recorded in many areas, including the fusiform gyrus and the inferior frontal gyrus (Barbeau et al., 2008). The second component peaks at 250 ms, followed by a third peak of inverse polarity (note that the polarity of the signal is arbitrary in SOBI, as it depends on the interaction between spatial and temporal components) and a maximum amplitude occurring between 400 and 600 ms (Figure 2a).

**Figure 1:**
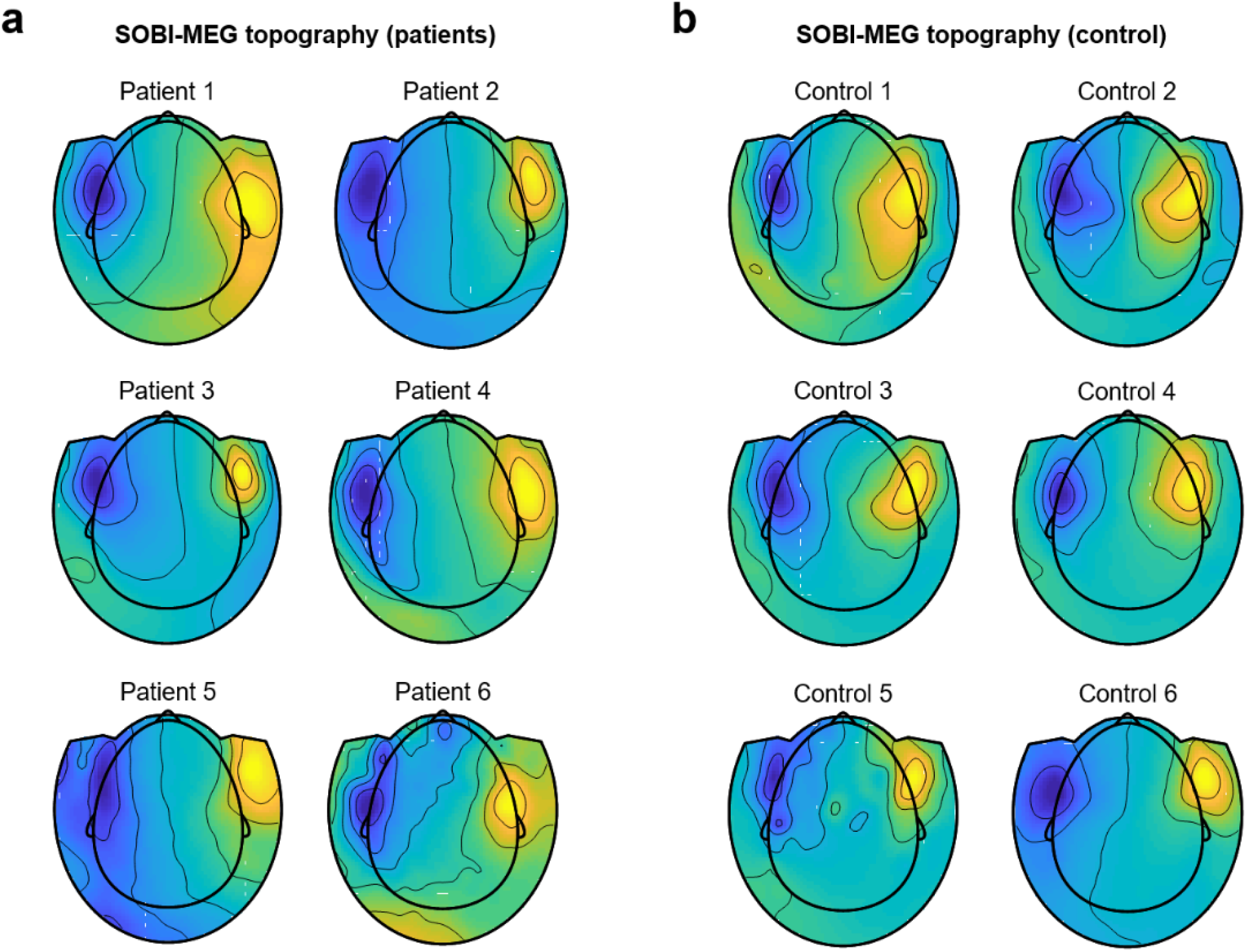
Second-Order Blind Identification (SOBI) magneto encephalographic (MEG) component of a putative deep source. a) Topography of the putative deep SOBI-MEG for all patients. The topography is extracted from the mixing matrixes obtained with the SOBI algorithm and represents the contribution of the SOBI source to each sensor. b) Topography of the SOBI-MEG in all controls.

**Figure 2:**
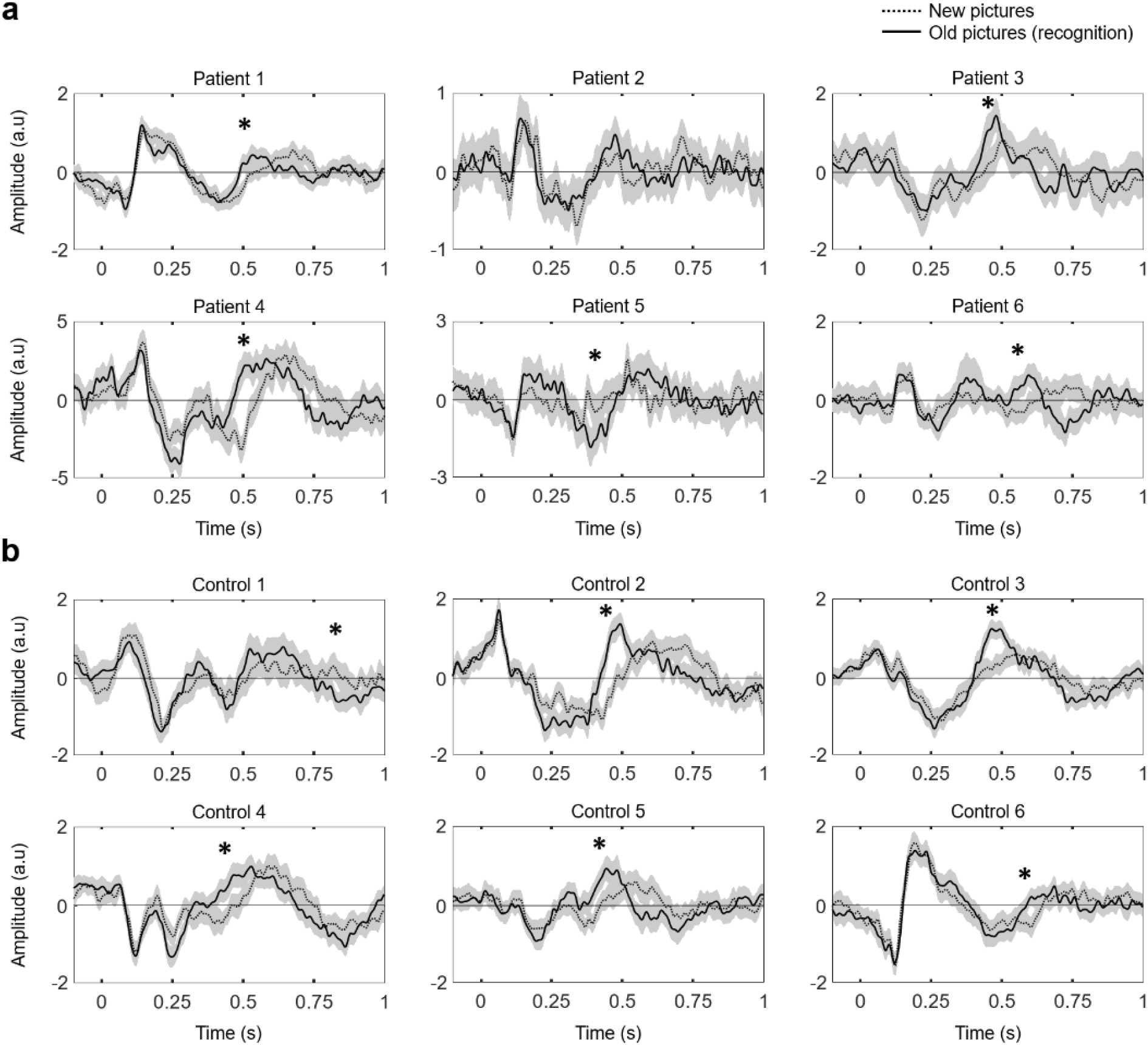
SOBI-MEG response to recognition. Response of the SOBI-MEG components represented in Figure 1. Solid and dashed traces are the averaged ERP (mean ± s.e.m. across trials) for old (recognition) and new trials, respectively. Stars indicate statistically significant differences in amplitude between old and new trials (unpaired t-test corrected using LFDR).

To confirm that the origin of the source was physiological and not pathological, we repeated the same memory protocol for MEG recordings obtained from six healthy volunteers. We identified a SOBI component with the same pattern as the one found in patients, sharing a similar deep topography (Figure 1b), and with the same time response, characterized by the maximums of opposite polarities at ~250 and ~500 ms (Figure 2b).

### SOBI-MEG differentiates old and new images

The hippocampal formation and surrounding areas have been shown to present characteristic ERPs in recognition memory tasks, during which their activity is modulated by object recognition (Axmacher et al., 2010; Barbeau et al., 2017, 2008; Merkow et al., 2015; Rugg and Curran, 2007). If the putative deep SOBI-MEG is related to the activity of mesial structures, a modulation of its ERP during the memory task is expected. To test this hypothesis, we compared the ERP responses in old versus new trials, testing for different amplitudes between both conditions at each time point. In good agreement with our hypothesis, we found statistically significant differences in five of the six patients and in all healthy controls (Figure 2). This modulation is observed at around 500 ms, with higher amplitudes, in absolute value, for old images. This result coincides with the hippocampal P600 responses for memory recognition (Barbeau et al., 2017, 2008; Merkow et al., 2015). Intriguingly, the differences between conditions are not only observed in amplitude but also in delay. ERPs have the same amplitude profile during the first 400 ms; then, there is an early peak in the recognition of old images (around 500 ms) that appears later in new images (around 600 ms, Figure 2). Together, these results support the deep origin of the SOBI-MEG topography.

### SOBI-MEG correlates with mesial areas in intracranial recordings

To further elucidate the putative origin of the SOBI-MEG component, we resorted to the intracranial field potentials recorded simultaneously with MEG from different regions. We compared the time courses of both datasets (SOBI-MEG and SEEG) by computing the zero-lag correlation, within each patient, between the SOBI-MEG topographies and all the recorded SEEG channels. A high correlation between SOBI-MEG and SEEG would indicate that both signals carry similar information, most likely because the source of the SOBI-MEG is close to the recorded channel. In Figure 3b, we have summarized the results for all patients, where color codes the correlation between the SOBI-MEG and the SEEG channels within the electrode (there were between 8 and 15 channels per electrode). For two patients (patients 4 and 6), the distribution of the correlation values and the threshold of significance with LFDR is represented in Figure 3c. The specific location of the contacts for each patient and their correlation with SOBI-MEG is represented in Figure 3d.

**Figure 3:**
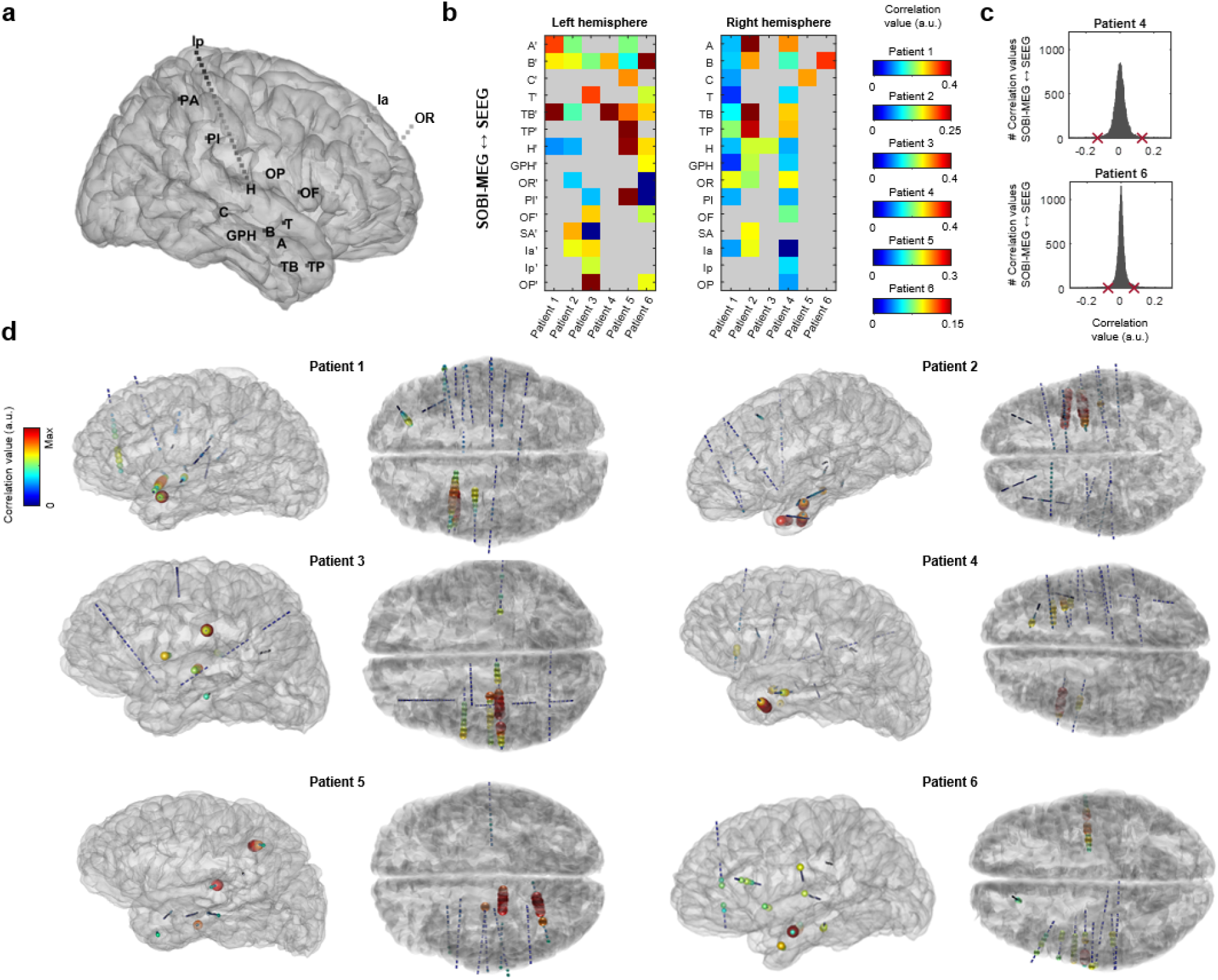
SOBI-MEG correlated with mesial intracerebral recordings. a) General intracerebral implantation scheme and nomenclature. b) Absolute value of zero-lag correlation between continuous time-series in the SOBI-MEG and in SEEG. On the y axis are the names of the SEEG electrodes. For each electrode, the channel with the highest correlation is represented. Light grey indicates that the electrode was not implanted in the patient. c) Distribution of correlation values between all SOBI-MEG and SEEG pairs for two patients. Red crosses are the threshold of significance obtained with LFDR. d) Reconstructed 3D brain mesh for each patient with SEEG contacts and their color-coded correlation with SOBI-MEG. Blue lines represent the contacts across each electrode, and both the color and size of the spheres indicate the correlation of that contact. Only significant correlation values are displayed.

In five out of six patients, the highest correlation values were found in electrodes located in mesial structures, maximal in TB (with contacts located in the anterior temporo-basal cortex, including the rhinal cortex), followed by B (with contacts located in the head of hippocampus), A (amygdala) and TP (temporal pole). Therefore, we focused the analysis on the brain structures recorded with these electrodes (Table 2). Note that electrodes TB, TP and A were not implanted in patient 3, limiting the opportunities for observing correlations with mesial structures. We also computed the contrast between experimental conditions (i.e., old vs new items) for each structure during the same window where the SOBI-MEG was modulated (400-600 ms). We found differences in the hippocampus, rhinal cortex and middle temporal gyrus of all patients with recordings in these locations. We also found differences in amygdala, temporal pole and inferior temporal sulcus in three out of the five patients implanted in these regions (Table 2).

**Table 2:** Correlation between SOBI-MEG and SEEG across mesial structures. A hashmark indicates that the region showed differences in the ERP for old and new images between 400 and 600 ms. A dash means that no SEEG contact was present in this region for this patient.

| Patient | Correlation |  |  |  |  |  |
| --- | --- | --- | --- | --- | --- | --- |
|  | 1 | 2 | 3 | 4 | 5 | 6 |
| *Hippocampus* | 0.24<sup>#</sup> | 0.10<sup>#</sup> | 0.10<sup>#</sup> | 0.24<sup>#</sup> | 0.12<sup>#</sup> | 0.09<sup>#</sup> |
| *Amygdala* | 0.28<sup>#</sup> | 0.10<sup>#</sup> | - | 0.19 | 0.14<sup>#</sup> | 0.01 |
| *Temporal pole* | 0.07 | 0.25<sup>#</sup> | - | 0.26 | 0.13<sup>#</sup> | 0.07<sup>#</sup> |
| *Rhinal cortex* | 0.34<sup>#</sup> | 0.18<sup>#</sup> | - | 0.40<sup>#</sup> | - | 0.10<sup>#</sup> |
| *Middle temp gyrus* | 0.19<sup>#</sup> | 0.18<sup>#</sup> | - | 0.29<sup>#</sup> | 0.20<sup>#</sup> | 0.06<sup>#</sup> |
| *Inferior temp gyrus* | 0.16 | 0.14 | - | 0.22 | 0.19 | - |
| *Inferior temp sulcus* | 0.38<sup>#</sup> | 0.20 | - | 0.37<sup>#</sup> | 0.05<sup>#</sup> | 0.04 |

The three mesial structures showing differences between old and new items in all the patients with available recordings were hippocampus, rhinal cortex, and middle temporal gyrus (Figure 4a). In figure 4b we show the averaged ERPs of the three regions for one patient. The hippocampus has a positive maximum at 400 ms in both old and new conditions, a component previously labelled hippocampal P600 (hP600, Barbeau et al., 2017). Moreover, this peak is followed by a fast decay, with a negative peak at 570 and 670 ms for old and new images, respectively. The rhinal cortex has a negative peak at 360 ms, that corresponds to the N360 previously reported in this area (Barbeau et al., 2017, 2008). Similar to the hippocampus, the negative peak is followed by a positive peak, with maxima at 510 and 650 ms for old and new images respectively. The middle temporal gyrus has the earliest response, with a double peak of opposite polarity at 150 and 210 ms. This activity may correspond to the visual response that can be recorded in several occipitotemporal brain areas (Barbeau et al., 2008). Moreover, following the dynamics of the other structures, it has a later response, with different delays for each condition. In short, the negative and positive peaks of old images were at 400 and 580 ms, respectively, while they were at 460 and 680 ms for new images.

**Figure 4:**
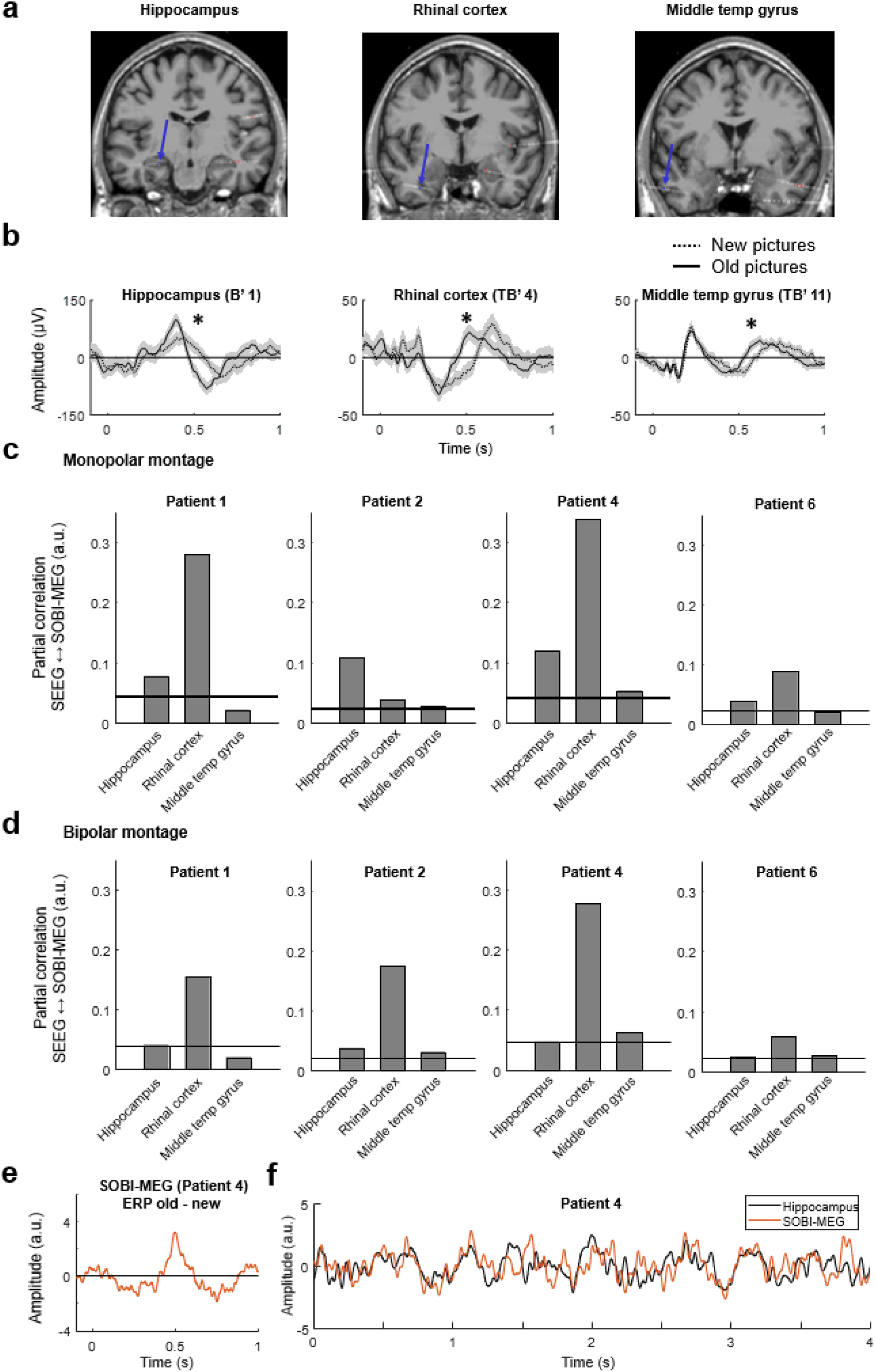
Partial correlation between SEEG signals and SOBI-MEG component. a) MRI (3D T1) with reconstruction of SEEG electrodes for patient 4. Arrows indicate the locations of the contacts for each selected region. b) Averaged ERPs for old (solid line) and new (dashed line) conditions from the three analyzed regions in patient 4 (mean ± s.e.m. across trials). Stars indicate statistically significant differences in amplitude between old and new trials (unpaired t-test corrected by LFDR). c) Absolute value of partial correlation between the SEEG recorded in three structures using a monopolar montage and the SOBI-MEG. Black lines represent the threshold of significance at p=0.025 for each patient (surrogate analysis). d) Same partial correlation analysis but using a bipolar montage for the SEEG recordings. e) Example of difference between averaged ERPs across trials in old minus new conditions in one patient. It can be appreciated the recognition effect at ~500 ms. f) Representative traces of hippocampal activity (monopolar SEEG montage) highly correlated with the SOBI-MEG component during the task.

The fact that the activity in these regions is correlated during the cognitive task (Figure 4b) may affect the correlation between SOBI-MEG and SEEG. A high correlation value may not indicate that we have recorded the actual brain source but another region whose activity is correlated with the source of interest. To differentiate these scenarios, we computed the partial correlation between the SOBI-MEG and the selected structures as recorded in SEEG (Figure 4c). This analysis tests whether the correlation between the SOBI-MEG and one region is direct (high partial correlation) or undirect, meaning that the common information is also in other regions (high correlation but low partial correlation). We excluded patients 3 and 5 from this analysis because the three regions were not recorded in those cases. The results confirmed the high partial correlation between the hippocampus and rhinal cortex with the SOBI-MEG. The highest partial correlation was found in the rhinal cortex (0.187 ± 0.145, mean ± SD), followed by the hippocampus (0.086 ± 0.037) and the middle temporal gyrus (0.031 ± 0.016). The values were significant in all patients for the hippocampus and rhinal cortex, but not the middle temporal gyrus (surrogate analysis, see Materials and methods).

One limitation of the monopolar montage used in this analysis is that activities arising from remote areas may affect the time course recorded at a given sensor due to volume conduction, and consequently the correlation between them and the SOBI-MEG. We converted the data with a bipolar montage, which aims to represent only the local activity, and we repeated the partial correlation analysis (Figure 4d) in the same sensors as in Figure 4c. The results corroborated the findings, with the highest partial correlation in the rhinal cortex in all patients (0.167 ± 0.090), followed by the hippocampus (0.037 ± 0.010) and the middle temporal gyrus (0.035 ± 0.019). The latter was significant in only three of the four analyzed patients (surrogate analysis). Finally, we focused on whether the memory effect in SOBI-MEG was the same as in the regions analyzed in Table 2. We measured the difference of the ERP between old and new conditions in both SOBI-MEG and SEEG contacts (Figure 4e) and we computed the correlation between them (Table 3). In good agreement with the previous results (Table 3 and Figures 4c and d), the maximum correlation value was found in the rhinal cortex in three out of four patients where we recorded this structure (0.668 ± 0.156), followed by middle temporal gyrus (0.584 ± 0.197) and hippocampus (0.534 ± 0.161). Overall, these results suggested that the origin of the SOBI-MEG is not a single structure, but it is the combination of multiple coactivated sources, including the hippocampus and the rhinal cortex (Figure 4f), confirming the notion of a mesial network.

**Table 3:** Correlation of the memory effect between SOBI-MEG and mesial structures. Bold values represent the area with highest correlation in each patient.

| Patient | Correlation ERP old-new (bipolar) |  |  |  |  |  |
| --- | --- | --- | --- | --- | --- | --- |
|  | 1 | 2 | 3 | 4 | 5 | 6 |
| *Hippocampus* | 0.56 | **0.60** | **0.61** | 0.74 | 0.27 | 0.45 |
| *Amygdala* | 0.61 | 0.53 | - | 0.83 | 0.34 | 0.28 |
| *Temporal pole* | 0.61 | 0.51 | - | 0.49 | 0.30 | 0.22 |
| *Rhinal cortex* | **0.69** | 0.55 | - | **0.88** | - | **0.55** |
| *Middle temp gyrus* | 0.67 | 0.40 | - | 0.84 | **0.64** | 0.37 |
| *Inferior temp gyrus* | 0.48 | 0.38 | - | 0.83 | 0.39 | - |
| *Inferior temp sulcus* | 0.16 | 0.46 | - | 0.76 | 0.45 | 0.53 |

### Source localization analysis places the origin of the SOBI-MEG in the mesial temporal lobe

Finally, we performed a source localization analysis on the SOBI topography. As there was no clear laterality in the SOBI-MEG topography, we assumed that the SOBI-MEG represented a bilateral activation of the mesial network (Piazza et al., 2020). Therefore, we computed the inverse problem with two dipoles that were symmetric with respect to the sagittal plane (Figure 5). We considered all points within the brain volume as potential solutions, without constraining the analysis to sub-cortical regions. The results revealed that the confidence interval for the source was not a single, well-localized point, but instead included a large area across the deep mesial cortex with a high goodness of fit (GOF>0.9; Figure 5). This reinforces the interpretation of the SOBI-MEG as a mesial network involved in memory recognition.

**Figure 5:**
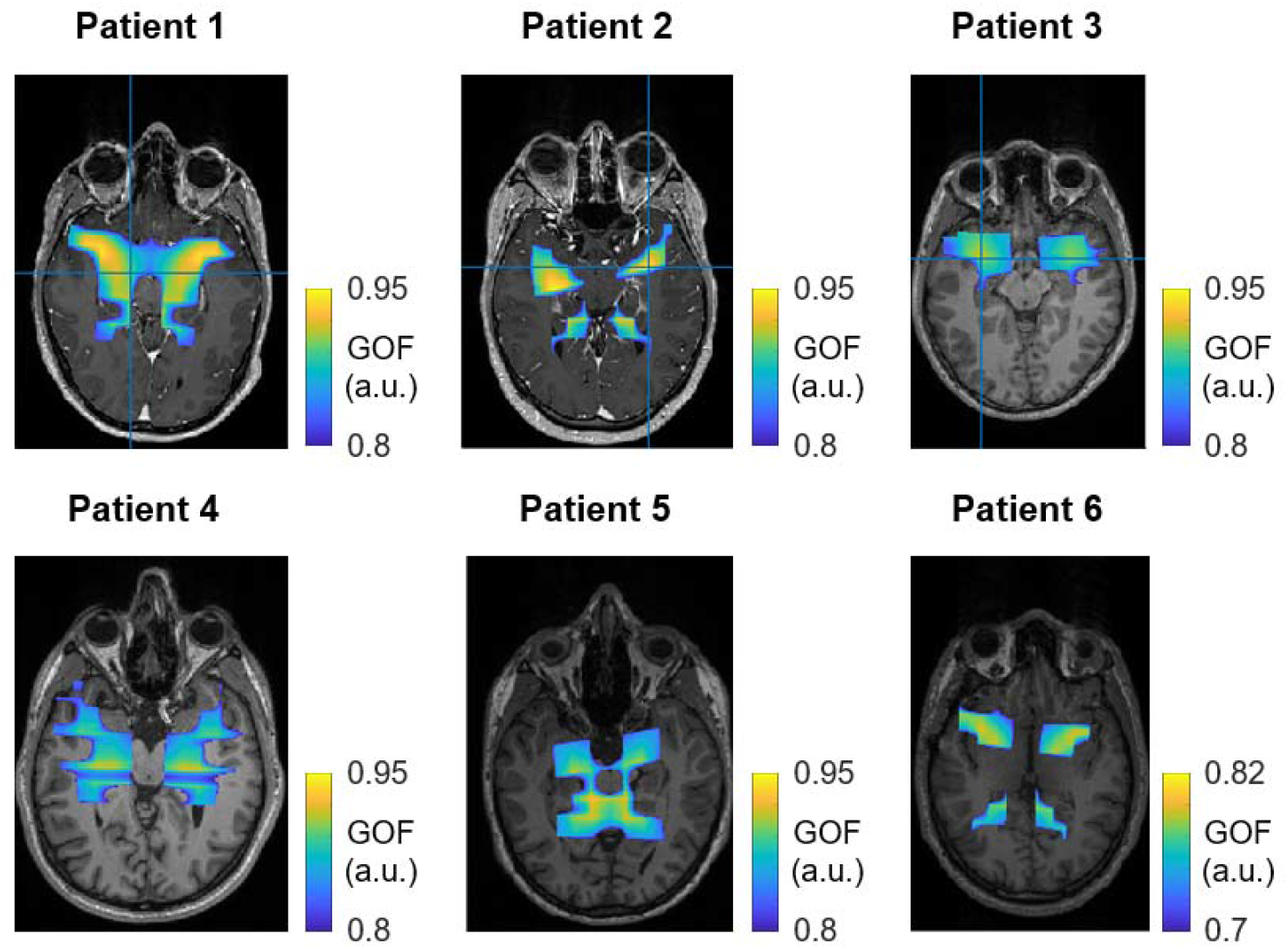
Source localization of the SOBI-MEG topography. Source localization of the SOBI-MEG topography with two symmetric dipoles for each patient.

## DISCUSSION

We report the identification of mesial activity in MEG during a recognition memory task in patients with focal drug-resistant epilepsy. Using SOBI, a BSS technique developed to disentangle the activity of sources that are mixed on the MEG sensors, we extracted one component, or SOBI-MEG, with a putative deep origin. This component was robust across patients, with similar topographies (Figure 1a) and time courses (Figure 2). It was likewise detected in all healthy controls. With simultaneous MEG-SEEG recordings serving as a ground truth of the brain activity, we confirmed the origin of the SOBI component as a mesial network comprising the hippocampus and the rhinal cortex.

The time-tested recognition memory task we used is well known to involve the mesial temporal lobe, with different responses between old and new images, especially in the hippocampus and the rhinal cortex (e.g. Barbeau et al., 2017, 2008; Despouy et al., 2020). We found that differences in the SOBI-MEG ERP were related in time to those observed in SEEG ERP across structures in the mesial temporal lobe. A partial correlation analysis confirmed the origin of the SOBI-MEG as a combination of the activities from, at least, the hippocampus and the rhinal cortex. Importantly, this analysis revealed that the SOBI-MEG is not only capturing the common activity between these structures, but also that it contains information specific to each node of the mesial network and not shared by the other areas included in the analysis (Figure 4c and d).

### Detectability of deep sources with BSS on simultaneous MEG-SEEG

The possibility to detect deep brain sources related to cognitive processes using surface sensors, with a special focus on the hippocampus, has been a matter of debates in recent years (Pu et al., 2018; Ruzich et al., 2019). Very few previous studies have used simultaneous intracranial and surface recordings to explore this issue. Such kind of co-registration is the only way to ensure the origin of the signals detected noninvasively, although it is not exempt of limitations (see below). Korczyn and colleagues (Korczyn et al., 2013) identified hippocampal theta on the MEG sensors, with a spatial pattern indicative of a deep source. While these results exhibited an accurate correlation between surface and deep signals, the authors limited their analysis to the hippocampus. Therefore, the possibility remains that the MEG sensor was recording a third region highly coherent at zero lag with the hippocampus rather than hippocampus itself. Moreover, because the activity recorded at the surface is the combination of several brain sources, the raw MEG data cannot be easily linked to a single source. It is necessary to resort to a methodology that can separate the different sources, such as BSS, to recover the time series associated with one region or one coherent network.

A popular BSS technique is independent component analysis (ICA). ICA assumes that the sources have specific spatial distributions that are invariant during the recording session (Comon and Jutten, 2010), and that the time-series of the components are independent, although it is robust to even high levels of source correlation (Makarova et al., 2011). Due to its versatility, ICA has been widely used to remove artifacts (Jung et al., 2000) and to separate neuronal sources in local field potentials (Herreras et al., 2015; Makarov et al., 2010), particularly in EEG and MEG (Debener et al., 2005; Delorme et al., 2002; Malinowska et al., 2014; Onton et al., 2005; Tang et al., 2002). Moreover, a recent study has compared the effectiveness of different BSS algorithms when analyzing dynamical functional connectivity on MEG (Tabbal et al., 2021). It has been argued that ICA relies not only on the independence of the sources, but also on their sparsity (Daubechies et al., 2009), which has been recently proposed as a key element to discriminate deep sources from the rest of the signals (Krishnaswamy et al., 2017). SOBI (Belouchrani et al., 1997, 1993), the alternative BSS method used here, is based on covariance; it minimizes the sum-squared cross-correlation between the components across a set of time delays and emphasizes processes that are correlated across neighboring time-points. We decided to use SOBI in this work because, contrary to ICA, it does not assume independency at each time point, and because its reliance on second order statistics makes it more robust to limited data length with a lower computational cost (Sahonero-Alvarez and Calderon, 2017). Furthermore, it has been argued that SOBI is more robust to variations in the underlying mixing matrix (Lio and Boulinguez, 2013) and in cases of temporal jitter (Huster et al., 2015).

ICA and simultaneous MEG-SEEG recordings were previously combined to identify epileptic spikes in the MEG sources (Pizzo et al., 2019). Because the signal of interest correlated with the hippocampus or the amygdala, but not with any other recorded structure, these structures were taken to be the sources of some independent components. However, it is important to note that the amplitude of the pathological epileptic spikes is much higher than the spontaneous physiological activity, and thus presumably much easier to detect from the surface. It remained unclear whether non-pathological hippocampal activity resulting from cognitive processes could be detected on MEG, which was the goal of the present study. The data reported here confirm that hippocampal and rhinal cortex activity can be detected with MEG during a cognitive task. Although the analyzed SOBI-MEG represents a network and not a single region, the partial correlation analysis revealed that both regions do contribute to the network and, importantly, that this contribution is unique and not shared by the other analyzed structures.

We further supported these results with a source localization analysis on the SOBI topographies (Figure 5). We decided to use two symmetric dipoles as brain sources obtained with BSS on MEG can often be explained with a dipole (Delorme et al., 2012), even though this is not guaranteed. Moreover, dipoles are justified for several reasons. First the non-dipolar part of the sources decreases very rapidly with distance and can be neglected for deep sources (Jerbi et al., 2004). Second, and in opposition with distributed source imaging methods, the equivalent dipole fitting does not absolutely require a noise covariance matrix, which cannot be easily estimated for BSS topographies. Third, the dipole fitting is robust to high level of correlations (Bénar et al., 2005) that are expected in cognitive paradigms. The source localization analysis was highly accurate to explain the SOBI-MEG topographies in Figure 1a, with large deep mesial areas with high GOF values (Figure 5). This is compatible with the interpretation of the SOBI-MEG as a mesial network including the hippocampus and rhinal cortex (Figure 4), but it is important to keep in mind that deep sources are expected to have large confidence intervals due to the low spatial frequencies on the scalp (large changes in position are needed to produce significant changes in the topography).

The challenge that still lies ahead is to fully reconstruct the time course of each structure in the MEG signal, and to differentiate it from other coactivated neuronal sources. The partial correlation analysis confirmed a certain degree of decorrelation between the sources conforming the SOBI-MEG, thus it is reasonable to expect that BSS algorithms can separate them. Still, the co-activation of the regions within the mesial network across trials can increase the temporal correlation and prevent perfect separation. Indeed, if there is not enough independency in space and in time between the sources, the BSS algorithm may not be able to separate them, and it will gather several sources into a single component. The combination of several protocols involving different structures may increase decorrelation and facilitate the separation of the regions.

### Memory response

The hippocampal formation, which comprises the hippocampus, the subiculum and the entorhinal cortex, is a key structure for spatial navigation and memory processes (Andersen et al., 2006; Eichenbaum, 1999; Morris et al., 1982). In humans, studies using SEEG have shown an increase of the P300 component in the ERP of the hippocampus during novelty detection (Halgren et al., 1995; Ludowig et al., 2010; Polich, 2007), while memory formation and face recognition elicited a hippocampal P600 component (Axmacher et al., 2010; Barbeau et al., 2017, 2008; Merkow et al., 2015; Rugg and Curran, 2007). In the rhinal cortex, an ERP with a double negative peak at N240 and N360 is also identified during memory recognition, concurrently with the activation of the amygdala and temporal pole (Barbeau et al., 2017, 2008; Despouy et al., 2020).

The fine analysis of the temporal dynamics of the SOBI-MEG during recognition memory is beyond the scope of the current study. Nevertheless, several features can be inferred from our results. In scalp EEG, the ERP is characterized by two components responding to old and new conditions (Hoppstädter et al., 2015; Rugg and Curran, 2007). The first occurs between 300-500 ms and is related to the prefrontal cortex. The second (400-800 ms) is linked with the hemodynamic response of the hippocampus and parahippocampal cortex (Hoppstädter et al., 2015), although this relation does not determine the origin of the electrical field in EEG. In the present work, the differences found in the SOBI-MEG occurred around 500 ms, corresponding probably to the second component reported in EEG. Intracranial signals corroborated the origin of the component, as old/new effects were also recorded in the hippocampus and rhinal cortex during the same time window (Figure 3b and 4a). Differences were observed not only in amplitude, but also in time, with shorter time delays for old versus new trials (Figure 2 and 4a, Trautner et al., 2004). Interestingly, the duration of this delay was between 100-150 ms, which corresponds to one theta cycle (~8 Hz). In the hippocampus, theta rhythms coordinate the activity between structures (Colgin et al., 2009; López-Madrona et al., 2020; López-Madrona and Canals, 2021), providing different temporal windows for communication, with different cells active at different phases (Schomburg et al., 2014). Furthermore, it has been suggested that both the encoding and the retrieval of events are segregated in the theta rhythm, either in the phase of the theta cycle (Hasselmo et al., 2002), or in different cycles of the same (Lopes-Dos-Santos et al., 2018; Zhang et al., 2019) or distinct theta oscillations (López-Madrona et al., 2020). To account for the old-new delay we observed here, we hypothesize that old and new images may be processed at distinct cycles of the hippocampal theta rhythm. For the old condition, the information about the memorized images in the thalamocortical working memory loop would coherently integrate in the hippocampus with the entorhinal input transmitting information about the external stimulus (de Vries et al., 2020; Raghavachari et al., 2006), thus facilitating the recognition in a first theta cycle. On the contrary, such integration would not occur in the new condition, as both inputs would mismatch. Consequently, the image would not be retrieved in the first theta cycle. The persistent external stimulus would enhance the entorhinal input in successive cycles, triggering the hippocampal circuit.

### Limitations of SEEG

There are two main restrictions inherent to SEEG recordings that may limit the conclusions of our study. Firstly, only a small part of the brain activity can be sampled with SEEG. Therefore, it is possible that the SOBI-MEG corresponds to a source not recorded with the available SEEG probes. Nonetheless, the timing of the different responses that we captured with SOBI-MEG have been reported across many studies primarily in the mesial structures included in the current study (Barbeau et al., 2017, 2008; Merkow et al., 2015). This reduces the likelihood that non-visible areas may have contributed to the SOBI-MEG component that we described. Secondly, the activity at each SEEG site is itself also composed of multiple field potentials converging on each contact, and the recorded activity may not be entirely local (Herreras, 2016). Bipolar montages (computed as the difference between adjacent contacts) allow a more precise characterization of local transient events. However, they may not recover the correct time-course of each area during ongoing field potentials (Fernández-Ruiz and Herreras, 2013; Martín-Vázquez et al., 2013) as, for instance, close generators with uncorrelated activities would cancel the resultant current. BSS methods have been proposed as a powerful methodology to extract the time course associated with different sources, either local or propagated, with promising results in the rat hippocampus and cortex (López-Madrona et al., 2020; Makarov et al., 2010; Ortuño et al., 2019). Nevertheless, the interpretation of the different components is not straightforward, and requires prior knowledge of the brain sources and the geometrical propagation of the fields in order to correctly infer the origin of the components (Herreras et al., 2015). More work is needed to investigate the use of BSS on SEEG signals in the context of cognitive paradigms.

## Conclusion

Blind-source separation methods reveal deep mesial activities in the MEG surface recordings whose localization could be reliably established through correlations with intracerebral SEEG recordings. This result has direct implications for clinical and cognitive neuroscience research. For example, accessing mesial structure activity with MEG could help to better understand processes underlying pathologies such as Alzheimer’s disease or epilepsy and, in particular, the detection of their early stages (Friston et al., 2015; Sharma and Nadkarni, 2020). The use of MEG could improve our knowledge of mesial temporal lobe processing in humans and of the implication of this region in memory processes. Overall, this opens new venues for the use of non-invasive MEG signals for characterizing physiological deep activity.

## ACKNOWLEDGMENTS

We thank Emmanuel Barbeau for his help in protocol design. This study was performed thanks to a FLAG ERA/HBP grant from Agence Nationale de la recherche, ANR-17-HBPR-0005 SCALES and UEFISCDI COFUND-FLAGERA II-SCALES, as well as thanks to the Swiss National Science Foundation (grants SNSF 192749 and CRSII5 170873 to S. Vulliemoz). The data was acquired on a platform member of France Life Imaging network (grant ANR-11-INBS-0006), supported in part by grants ANR-16-CONV-0002 (ILCB) and the Excellence Initiative of Aix-Marseille University (A*MIDEX).

## DATA AVAILABILITY STATEMENT

The original raw data supporting the findings of this study are available upon request to the corresponding authors.

